# Blood pressure lowering enhances cerebrospinal fluid efflux primarily via the lymphatic vasculature

**DOI:** 10.1101/2023.07.11.548482

**Authors:** Jari Jukkola, Mika Kaakinen, Abhishek Singh, Sadegh Moradi, Hany Ferdinando, Teemu Myllylä, Vesa Kiviniemi, Lauri Eklund

## Abstract

**Background:** Inside the incompressible cranium, the volume of cerebrospinal fluid (CSF) is directly linked to blood volume: a change in either will induce a compensatory change in the other. Vasodilatory lowering of blood pressure has been shown to result in an increase of intracranial pressure, which, in normal circumstances should return to equilibrium by increased fluid efflux. In this study, we investigated the effect of blood pressure lowering (BPL) on fluorescent CSF tracer absorption into the systemic blood circulation.

**Methods:** BPL was performed by an i.v. administration of nitric oxide donor sodium nitroprusside (5 µg kg^-1^ min^-1^) or the Ca^2+^-channel blocker nicardipine hydrochloride (0.5 µg kg^-1^ min^-1^) for 10 and 15 to 40 mins, respectively. The effect of BPL on CSF clearance was investigated by measuring the efflux of fluorescent tracers (40 kDa FITC-dextran, 45 kDa Texas Red-conjugated ovalbumin) into blood and deep cervical lymph nodes.

**Results:** Nicardipine and sodium nitroprusside reduced blood pressure by 32.0 ± 19.6% and 22.0 ± 2.5%, while temporarily elevating in intracranial pressure by 14.0 ± 6.0% and 11.6 ± 2.0%, respectively. BPL significantly increased tracer accumulation into deep cervical lymph nodes and systemic circulation, but reduced perivascular inflow along penetrating arteries in the brain. The enhanced tracer efflux by BPL into the systemic circulation was markedly reduced (-66.7%) by ligation of lymphatic vessels draining into deep cervical lymph nodes.

**Conclusions:** This is the first study showing that CSF clearance can be improved with acute hypotensive treatment and that the effect of the treatment is reduced by ligation of a lymphatic drainage pathway. Enhanced CSF clearance by BPL may have therapeutic potential in diseases with dysregulated CSF flow.

## Background

Cerebrospinal fluid (CSF) serves as a protective fluid for the brain and spinal cord, it maintains metabolic homeostasis and functions as a waste sink for the central nervous system (CNS) derived metabolic waste products (Flexner, 1933; Iliff et al., 2013a; Simon & Iliff, 2016). CSF is not only confined to the superficial parts of the CNS, but it also extends deeper into brain parenchyma along the perivascular spaces of penetrating arteries, mixes with brain interstitial fluid and thus may participate in the clearance of brain metabolic waste products in the absence of parenchymal lymphatic vessels (Kress et al., 2014; Xie et al., 2013). From the CSF, macromolecules can be directly transferred to the bloodstream via venous sinuses (Clark, 1920; Jiang et al., 2021; Potts & Deonarine, 1973; Zakharov et al., 2004) or absorbed into dural tissue and meningeal lymphatic vessels distributed along venous sinuses in the dorsal and basal skull (Ahn et al., 2019; Antila et al., 2017a; Aspelund et al., 2015; das Neves et al., 2021; Louveau et al., 2015). Other exit pathways include the nasal lymphatic route and drainage along cranial nerves (Brady et al., 2020; Hoffmann et al., 2019; Johnston et al., 2004; Mollanji et al., 2002; Silver et al., 2002; Turliuc et al., 2016; Wood et al., 2019). Many recent findings imply the essential role of the meningeal and nasal lymphatic systems (Brady et al., 2020; Da Mesquita et al., 2021; Louveau et al., 2017; Mollanji et al., 2002; Pedler et al., 2021; Proulx, 2021; Silver et al., 2002; Turliuc et al., 2016), e.g., enhancement of meningeal lymphatic function with VEGF-C treatment was recently demonstrated to improve cognitive behavior and facilitate the drainage of CSF into deep cervical lymph nodes in aged mice (da Mesquita et al., 2018; Hsu et al., 2021).

While the relative contribution of the proposed clearance pathways in CSF clearance and underlying mechanistic principles still await further detailing, CSF production is decreased during aging and disrupted CSF flow is associated with cognitive decline in Alzheimer’s disease (AD) (Attier-Zmudka et al., 2019; Fleischman et al., 2012; Harrison et al., 2020; May et al., 1990; Rubenstein, 1998; Stoquart-ElSankari et al., 2007). In addition, the presence of biomarkers of AD in CSF samples has relevance in disease progression predictions (Lee et al., 2020; Wolfsgruber et al., 2017). Altogether, these findings indicate the importance of adequate CSF production, circulation, and absorption in brain homeostasis, whereas aging and diminished CSF drainage predispose accumulation of harmful metabolites due to attenuated CSF dynamics. Furthermore, increased blood pressure (BP) reduces CSF convection (Mestre et al., 2018) and conversely tight control of blood pressure can reduce the risk of mild cognitive impairment (Williamson et al., 2019). Clearly, enhancement of CSF circulation and clearance deserves further investigation.

In this study, we reasoned that modulation of brain pressure-volume balance by lowering systemic blood pressure alters CSF absorption dynamics according to the Monro-Kellie doctrine, which describes the pressure-volume relationship of brain components inside a rigid skull (Kellie, 1824; Wilson, 2016). We studied the drainage efficiency of 40 kDa FITC-dextran (FD40) and 45 kDa Texas Red-conjugated ovalbumin injected into the cisterna magna (CM) with acute reduction of blood pressure by intravenous (i.v.) infusions of vasodilators. We observed that acute blood pressure lowering (BPL) improved CSF absorption into the systemic circulation and that absorption was markedly reduced by ligation of intact lymphatic drainage pathway.

## Results

### Blood pressure lowering (BPL) temporarily increases intracranial pressure and enhances tracer efflux into blood

An i.v. injection of nicardipine resulted in a continuous decline in BP (Fig. 1A) starting 8 min of infusion and reaching the lowest value at 40 min (68.3 ± 19.6% of the baseline). In ∼15 min, BP dropped by -19.0 ± 11.5% while intracranial pressure (ICP) increased significantly (+14.0 ± 7.0%) and returned to baseline levels within ∼40 min (Fig. 1C).

**Figure 1.**
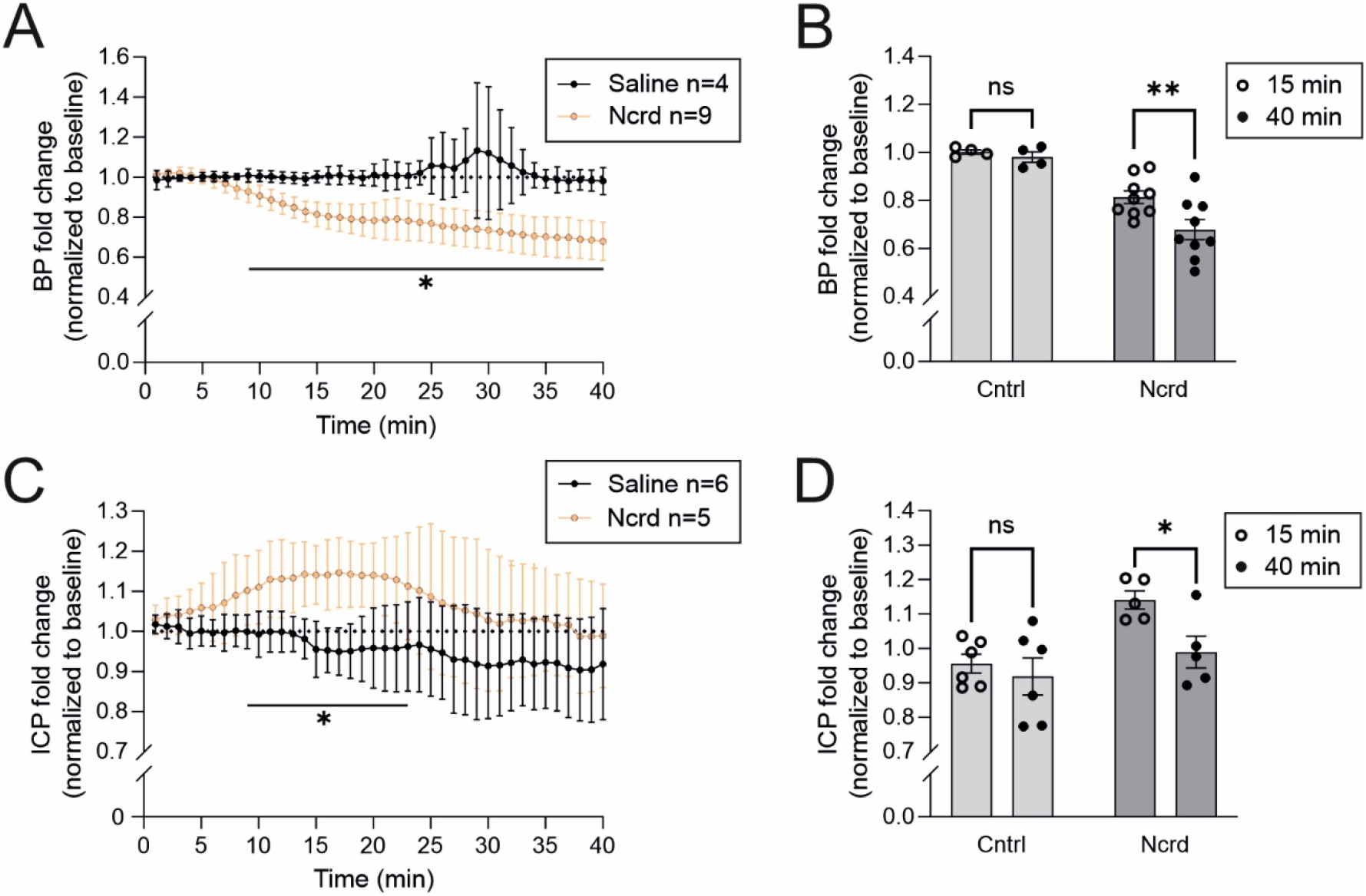
Cerebral blood and intracranial pressures during BPL. **A, C,** Blood pressure (BP) and intracranial pressure (ICP) during BPL with nicardipine (Ncrd) on separate groups of mice. Multiple unpaired t-tests with Welch’s corrections. **B, D,** Comparisons of selected timepoints in control and BPL groups. Ordinary two-way ANOVA with Šidák’s multiple comparisons. **A – D,** \**P* < 0.05, ns = non-significant. **A, B,** n = 4 – 9 mice per group. **C, D,** n = 5 – 6 mice per group. Filled and open circles in B and D represent individual mice at different timepoints taken from the graphs A and C, respectively.

To investigate if the increased ICP improves the drainage of macromolecules into the systemic blood circulation, we injected 4 µl of FD40 into the CM, followed by blood sampling before and at the end of BPL treatment or saline infusion (Fig. 2A). FD40 concentration increased 2.0-fold (3.0 ± 1.5 µg ml^-1^ vs 6.0 ± 1.0 µg ml^-1^) after the 15-min BPL treatment (Fig. 2B) and a 2.7-fold (4.5 ± 1.5 µg ml^-1^ vs 12.4 ± 7.5 µg ml^-1^) after the 40-min BPL treatment (Fig. 2C). No significant change in FD40 concentration was found in controls receiving only saline (Fig. 2D).

**Figure 2.**
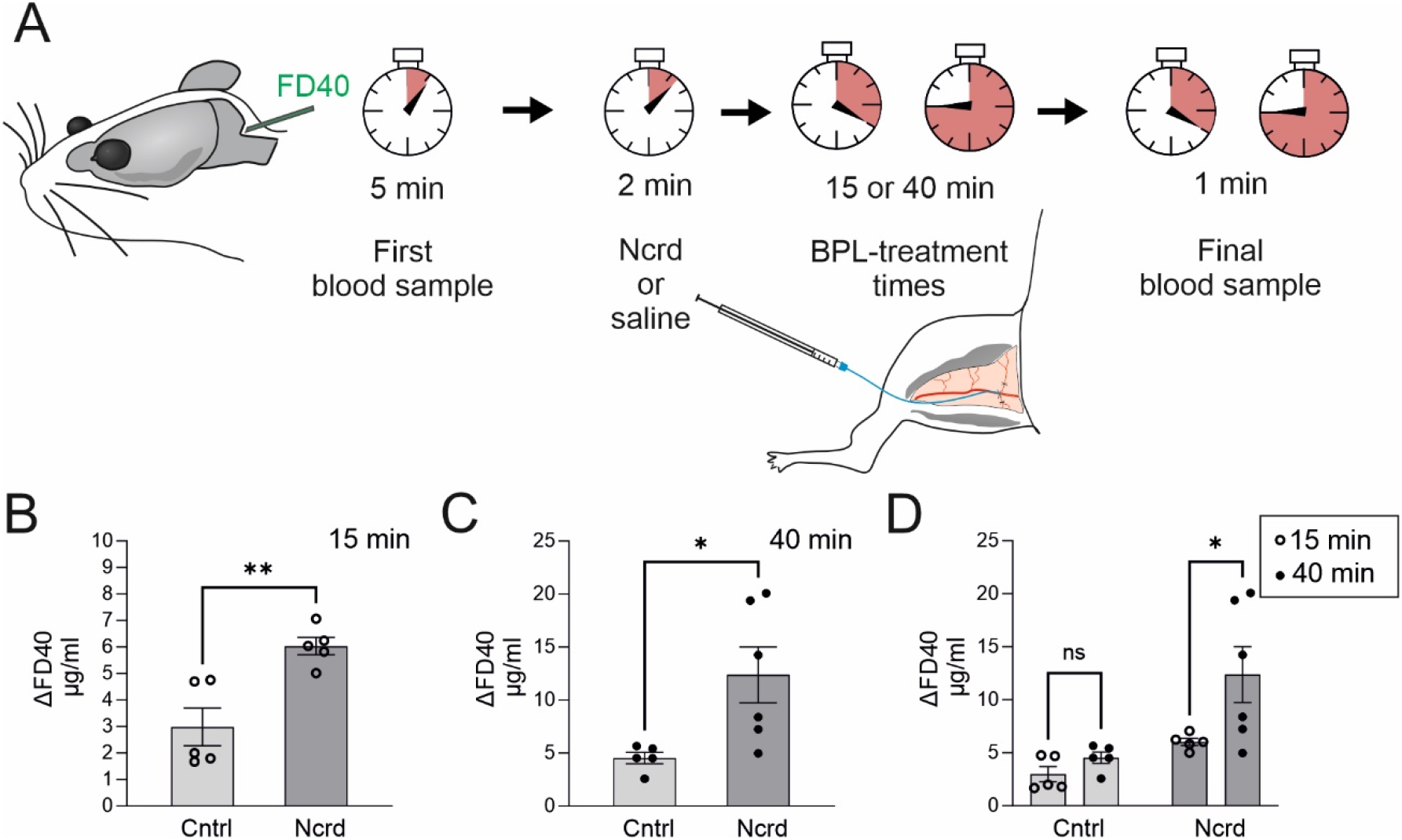
Manipulation of CSF tracer efflux with BPL. **A**, Experimental scheme and timeline to investigate the effect of BPL on CSF tracer efflux. 40 kDa FITC-dextran (FD40) injection into the cisterna magna was followed by blood sampling from femoral vein 5 minutes later, with an i.v. (femoral vein) infusion of nicardipine (Ncrd) or saline (Cntrl) starting 2 minutes after the first blood sampling for either 15 or 40 min, after which the final blood sampling was performed. **B, C,** FD40 concentration change in blood during 15-min and 40-min BPL, Mann-Whitney *U* test and unpaired t-test with Welch’s correction, respectively. **D,** Comparison of selected timepoints in control and BPL groups. Two-way ANOVA with Šidák’s multiple comparisons. **B – D,** \**P* < 0.05, ns = non-significant. n= 5 – 6 mice per group. Filled and open circles in B-D represent individual mice.

### Blood pressure lowering increases tracer efflux into deep cervical lymph nodes but reduces perivascular influx in the brain

We next investigated if BPL increases tracer drainage into the deep cervical lymph nodes (dcLNs), the major LNs draining the meningeal lymphatics (Antila et al., 2017b; Aspelund et al., 2015; Bradbury et al., 1981; Louveau et al., 2015; Maloveska et al., 2018; Song et al., 2020). Additionally, to understand the CSF movement in brain, we investigated perivascular (glymphatic) influx in brain sections (Iliff et al., 2013b; Keil et al., 2022). We injected fixable Texas Red-conjugated ovalbumin with a molecular weight similar to FD40 into the CM. We found that perivascular tracer penetration was significantly decreased in the brain sections (14.1 ± 7.0 a.u vs. 3.1 ± 1.5 a.u.) and tracer content in dcLNs was significantly increased (40.9 ± 30.6 a.u. vs 115.4 ± 75.1 a.u.) in BPL-treated mice when compared to controls receiving only saline (Fig. 3).

**Figure 3.**
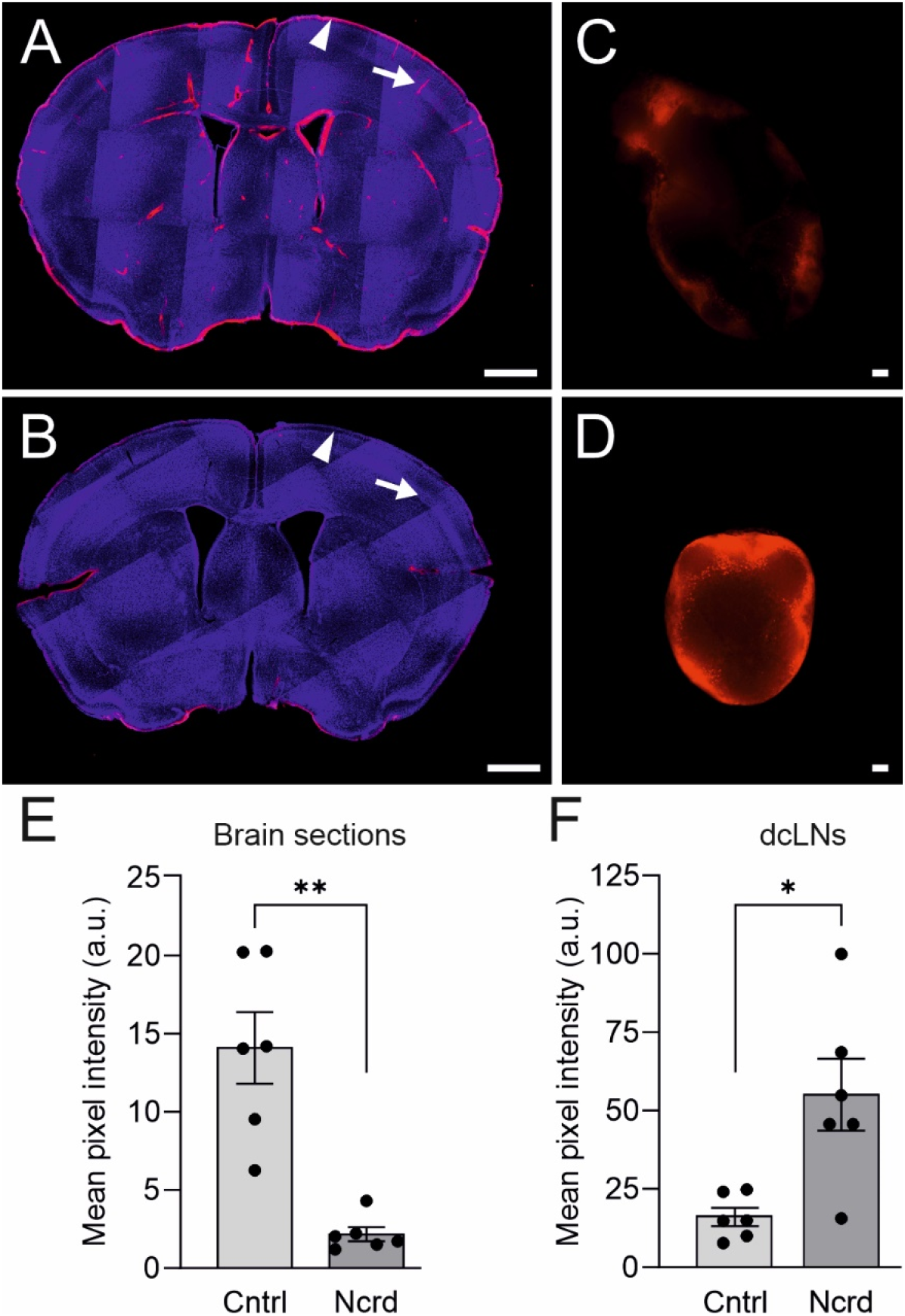
Perivascular brain influx and drainage of Texas Red-conjugated ovalbumin into dcLNs. **A**, A brain section from a control mouse showing deep parenchymal penetration (arrow) and cortically located 45 kDa Texas Red-conjugated ovalbumin (arrowhead). **B,** A brain section of a mouse after BPL displays significantly reduced presence of tracer, as quantified in E. **C, D,** Representative images demonstrating that BPL (D) improves tracer uptake into to dcLNs compared to saline-treated controls (C). **A – D,** Blue, DAPI; Red, 45 kDa Texas Red-conjugated ovalbumin. **E, F,** Fluorescence intensity quantitation from serial brain sections and dcLNs, \**P* < 0.05, unpaired t-test with Welch’s correction and unpaired t-test, respectively. Scale bars for brain sections and dcLNs are 1 mm and 100 µm, respectively. **A – F,** n = 6 mice per group. Filled and open circles in E and F represent individual mice.

### Blocking deep cervical lymphatic vessels attenuates tracer outflow during blood pressure lowering

To test the contribution of extracranial lymphatic drainage via dcLNs to the BPL effect on CNS clearance, we performed ligation of afferent lymphatic vessels (Fig. 4) that drain into dcLNs and quantified the FD40 concentration from the blood. Lymphatic efflux has been previously shown to precede blood accumulation which occurs 15-30 min after tracer injection into the CSF (Ma et al., 2017). An i.v. injection of rapid-acting sodium nitroprusside (SNP) resulted in a drop in BP (22.0 ± 2.5%) and concomitant elevation of ICP (11.6 ± 2.0%) (Figs. 4E and F) while the ligation alone did not significantly elevate ICP within the estimated time window in which the separate efflux experiments were performed (Fig. 4G). Ligation of afferent lymphatic vessels abolished tracer drainage into dcLNs during BPL with SNP, resulting in a significant reduction of FD40 in blood (5.1 ± 1.5 µg ml^-1^ vs 1.7 ± 2.2 µg ml^-1^), thus suggesting that BPL-induced improvement of tracer absorption from the CSF is primarily through the lymphatic vasculature (Fig. 4H).

**Figure 4.**
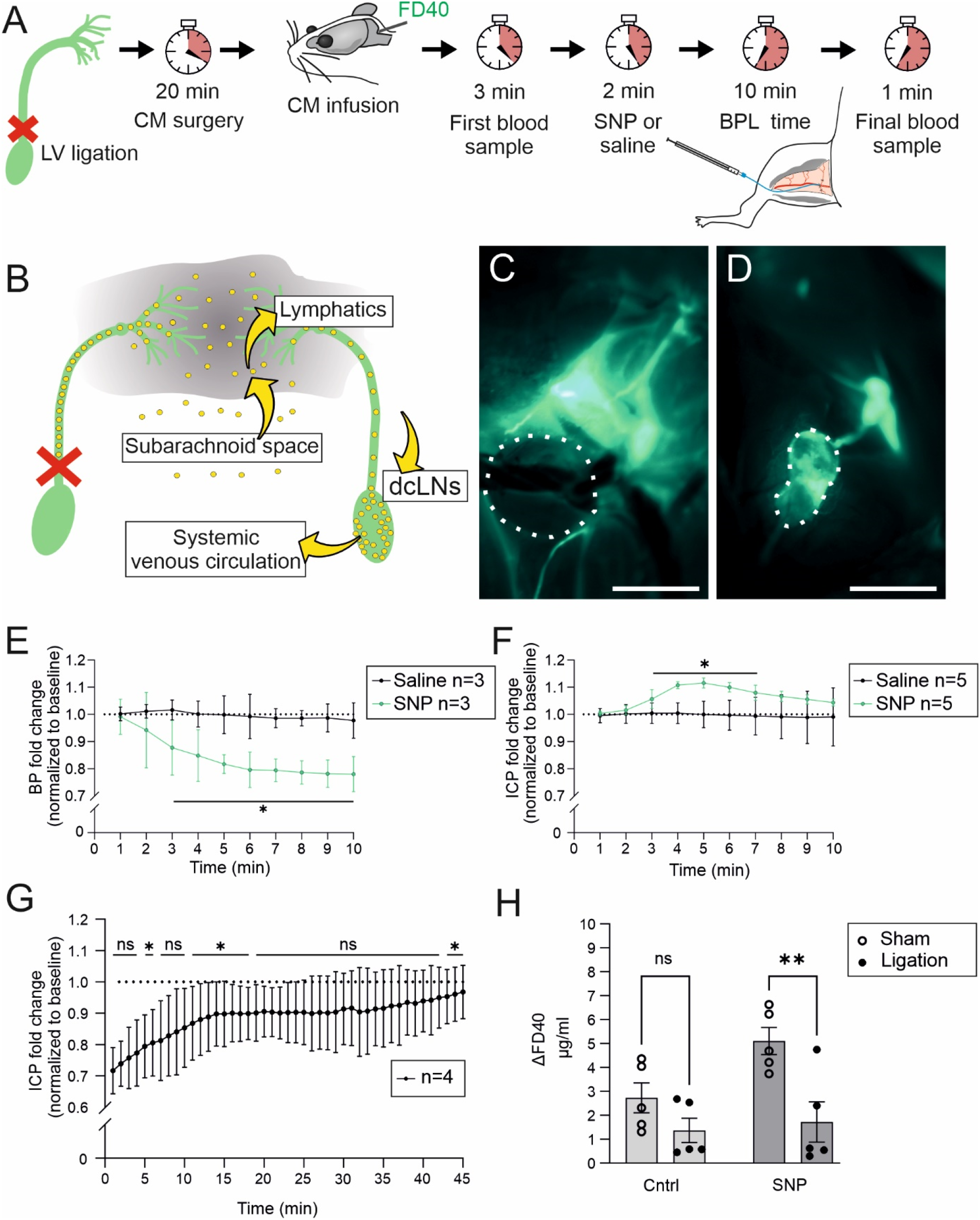
CSF tracer drainage via extracranial lymphatics to systemic blood circulation during BPL. **A**, Experimental scheme and timeline of afferent lymphatic vessel ligation, FD40 injection and blood sampling. CM infusion of FD40 began ∼20 min after lymphatic vessel (LV) ligation, followed with first blood sampling (3 minutes after the infusion), 10-min BPL or saline treatment (2 minutes later) and final blood sampling. **B,** An illustration of afferent LV ligation (crossed red lines) which obstructs tracer uptake (yellow spheres) into a lymphatic efflux pathway (yellow arrows) and deep cervical lymph nodes (dcLNs). **C,** Drainage of FD40 into a dcLN (white dashed lines) via the afferent lymphatic vessel is abolished after ligation. **D,** FD40 drainage into a dcLN on the sham-operated side of the same animal as in C. **E, F,** Blood-(BP) and intracranial pressure (ICP) measurements during BPL with sodium nitroprusside (SNP), multiple unpaired t-tests with Welch’s correction. **G,** ICP after ligation of dcLNs, one-way ANOVA with repeated measurements and Dunnett’s multiple comparisons. **H,** Concentration changes of FD40 in blood after BPL with SNP in sham-operated and ligated groups, two-way ANOVA with Šidák’s multiple comparisons. **C, D,** Scale bar, 1 mm. **E–H,** \**P* < 0.05, ns = non-significant. n = 3 mice (E), n = 5 mice (F), n = 4 (G) and n = 5 (H). Filled and open circles represent individual mice.

## Discussion

The major finding in this study is that acute BPL increases tracer efflux from the CSF to the lymphatic vessels that drain into the venous system through the collecting ducts. CSF clearance is highly relevant for brain homeostasis, for the recently proposed brain clearance mechanisms and in the pathogenesis of neurodegenerative diseases. However, there are no current treatment options to augment the clearance function. Our study presents novel method to enhance cerebral CSF flow in mice, suggesting that cerebral macromolecular clearance can be increased by a clinically feasible treatment.

As stated by the Monro-Kellie doctrine, a change in the volume of one compartment in the cranium must follow a compensatory volume displacement in others (Kellie, 1824). Vasodilation can increase vascular compliance while decreasing the compliance of the intracranial space, suggesting that even small changes in blood volume can affect ICP (Kim et al., 2009). Accordingly, BPL resulted in an increase in ICP, a common side effect reported also in clinical studies (Anile et al., 1981; Cottrell et al., 1978; Levy, 2005; Nishikawa et al., 1986). Increased ICP coincides with the CSF volume displacement into blood as seen as increased tracer efflux. Contrasting with the diminishing effect of increased ICP on CSF inflow into brain via perivascular spaces, intraperitoneal injection of hyperosmotic mannitol reduces ICP and increases perivascular CSF inflow in mice (Plog et al., 2018). The pathological and sustained rise in ICP is commonly observed in traumatic brain injury (Balestreri et al., 2006). However, in contrast to an acute (∼15 min with nicardipine, ∼4 min with SNP) and a rather mild (∼11 – 14%) elevation in ICP we observed, a long-lasting (> 2 h) with more than two-fold elevation of ICP may compromise CSF drainage via the meningeal lymphatics (Bolte et al., 2020; Xiang et al., 2022).

The accumulation of Texas Red-conjugated ovalbumin (45 kDa) into dcLNs was evident in ∼40 min following injection at CM and was significantly facilitated by BPL-treatment. Furthermore, ligation of afferent lymphatic vessels draining to dcLNs significantly reduced the blood uptake of FD40 within ∼18 min following injection into CM. In a previous study, tracers injected into the CSF exhibited a 25 to 30 min time delay before appearing in the bloodstream, whereas they were present in draining lymph nodes within 15 min (Ma et al., 2017). Consistent with this finding, mice lacking a meningeal lymphatic system showed delayed clearance of extracellular tau-protein into blood (Patel et al., 2019). A direct lymph transport is thus likely the main route for CSF absorption preceding a significant transit to the blood. Blood vessels may take up molecules from CSF parallelly with lymphatic drainage but act slower and less efficiently.

Increased tracer efflux from the CSF coincided with reduced tracer inflow into the brain. This could be due to reduced amount of available tracer for paravascular entry along penetrating arteries as shown previously in awake mice (Ma et al., 2019). It is also possible that vasodilation of pial- and penetrating arteries reduces the perivascular space, diminishing directional fluid flow. It was indeed shown recently that vasodilation by whisker stimulation reduces periarterial space and CSF flow velocity, but whisker stimulations and/or optogenetically induced vasoconstrictions in sequence lead to cumulative fluid inflow volume (Holstein-Rønsbo et al., 2023). Thus, constantly altered vascular tone rather than chronic change (as in the present study) might be prerequisite for increased perivascular inflow as also discussed in the aforementioned article by Holstein-Rønsbo *et al*. (2023). With vasoactive agents, variances in dose-response relationship may contribute on the net direction of perivascular flow. For example, dexmedetomidine, a selective alpha 2-adrenoceptor agonist, reduces systemic blood pressure, cerebral blood flow and ICP (in laparoscopic surgery) while facilitating brain delivery of intrathecally administered drugs (Drummond et al., 2008; Lilius et al., 2019; Sahay et al., 2018). However, dexmedetomidine has also dose-dependent vasoconstrictive effects in cerebral arterioles (Asano et al., 1997) which may explain improved CSF inflow apart from the systemic BPL-effect. In support of this, stroke associated arterial vasoconstriction enlarges the perivascular space and leads to increased CSF inflow into the brain (Mestre et al., 2020). Intracranial vasoconstriction has also been reported for nicardipine treatments in some studies (Lahiri et al., 2018), while in others nicardipine either increases or leaves cerebral blood flow unaltered and causes vasodilation (Gaab et al., 1985; Nagahama et al., 1997; Takenaka & Handa, 1979). For example, an i.v. dosage of 1 to 30 µg kg^-1^ increases cerebral blood flow and ICP in monkeys (Takenaka & Handa, 1979). In the present study, the observed increase in ICP likely implies a volume increase in the cerebral vasculature similar to a previous study (Takenaka & Handa, 1979). It is thus likely that BPL-induced cerebral vasodilation disrupts the glymphatic intake by reducing perivascular flow, mixing of CSF and interstitial fluid and the proposed convective flow through the interstitium (Iliff et al., 2012).

A recent finding shows that vasomotion is a motive force driving nanoparticles from brain alongside arteries (van Veluw et al., 2020), suggesting that brain metabolites are cleared along the vasomotion-driven intra-mural peri-arterial (IPAD) clearance pathway (Aldea et al., 2019). Interestingly, in awake mice, regional vasodilation by local functional hyperemia significantly increases the clearance rate of extravasated fluorescence tracers from the brain parenchyma (van Veluw et al., 2020). During regional activation-induced hyperemia, cardiovascular pulsation also increases locally, potentially driving perivascular flow in humans (Huotari et al., 2022). In accordance, vasoactive drugs have been shown to improve parenchymal clearance of harmful metabolites, provide neuroprotection, and alleviate hypertension-associated reduced cerebral blood flow (Maki et al., 2014; Tryambake et al., 2013). For example, cilostazol (a phosphodiesterase III inhibitor) that has multiple effects on the vasculature, was shown to facilitate the perivascular drainage of amyloid beta in Tg-SwDI mice, plausibly by increasing pulse duration time and elasticity of vascular SMCs (Maki et al., 2014). In the SMCs, cilostazol stimulates interleukin-1β-induced nitric oxide (NO) production (Ikeda et al., 1996; Kitade et al., 1996), which may explain its vasoactive properties (Yasuda et al., 1985). NO donors, including SNP and sodium nitroglycerin, are used in clinics for the treatment of cardiovascular diseases, but they have also been shown to downregulate the N-methyl-D-aspartate (NMDA) receptors (Choi et al., 2000), in which overactivity can impair cognitive functions in mice (Li et al., 2022) and hasten AD progression in humans (Bukke et al., 2020; Greenamyre et al., 1988; Paoletti et al., 2013; Yeung et al., 2021). However, due to the systemic vasodilatory action, long-term usage of these drugs increases the risk of hypotension. Recent progress in drug development has produced NO donors which specifically release NO in brain tissue (Liu et al., 2017). A nitrate moiety coupled to memantine, a non-competitive NMDA receptor antagonist, which is used to treat moderate to severe AD, enables brain tissue specific NO release with consequent cerebral vasodilation and increased cerebral blood flow (Luo et al., 2019). In this current study, BPL resulted in systemic vasodilation that facilitates clearance of macromolecules from CSF. It is thus plausible that controlled vasodilation would contribute to CSF-mediated metabolite clearance from parenchyma to extracranial lymphatics and ultimately to systemic circulation.

## Conclusion

In conclusion, our data show that CSF dynamics can be manipulated by BPL-treatment, which provides empirical justification for the Monro-Kellie doctrine in a non-pathological context. Moderate increase in ICP by BPL resulted in enhanced filtration of tracers into the bloodstream primarily through extracranial lymphatics, whereas direct transfer to the bloodstream seems to play a less significant role during the BPL-treatment in mice. Further investigations addressing the translational and therapeutic potential regarding the temporary enhancement of CSF absorption and lymphatic macromolecular drainage with BPL are warranted.

## Methods and materials

### Animals and housing

2- to 3-month-old C57/Bl6N (Charles River) female mice were used in all experiments. All animals were housed in temperature- and humidity-controlled rooms with a 12h light/12h dark cycle. Teklad Global Rodent diet (Harlan Teklad, USA) and untreated tap water were offered *ad libitum*. Mice were anaesthetized with subcutaneous injection of ketamine (75 mg kg^-1^ or 100 mg kg^-1^) and xylazine (10 mg kg^-1^) (KX). Lidocaine (10 mg kg^-1^) was administered subcutaneously to surgical sites 5 min before operations.

### Cisterna magna (CM) cannulation and tracer infusion

After presurgical preparations the mice were connected to a stereotaxic device. After checking surgical site pain reactivity with forceps, neck muscles were gently separated to expose the CM. A dental needle (30G) connected to a polyethylene tube (BTPE-10) filled with saline was used to cannulate the CM as described previously (Xavier et al., 2018). FITC-dextran 40 kDa (FD40) (Sigma, FD40S-250MG; 25 mg ml^-1^) or 45 kDa Texas Red-conjugated ovalbumin (Thermo Fisher Scientific, O23021; 1 mg ml^-1^) were diluted in saline and withdrawn via a connecting line into the cannula. Unlike FD40, Texas Red-conjugated ovalbumin could be fixed in tissues, allowing *ex vivo* quantitation. 4 µl of tracers were injected with a rate of 2 µl min^-1^ into the CM by a Hamilton syringe with a 30-gauge needle filled with saline and operated with a microinjector (KD Scientific model LEGATONANO). After infusion, excess cannula was sealed by using an electrocauter (Bovie, change-a-tip DEL1). An infrared lamp or a heat pad (set to +37°C) was used to help maintain the body temperature. The first blood sample was collected within ∼5 min after the infusion, and the final sampling ∼18 min (dcLN ligation experiments) or ∼45 min later in the other experiments. Blood sampling was from the femoral vein (∼160 µl for the 1^st^ sample) and carotid artery (∼180µl – 200µl for the 2^nd^ sample) and mice were sacrificed immediately after collecting the second blood sample.

### Blood and ICP pressure measurements

A BTPE-50 polyvinyl cannula (filled with 0.9 % NaCl supplemented with 500 IU ml^-1^ heparin) was inserted into the permanently ligated left external carotid artery to access the common carotid artery as previously described (Elamaa et al., 2018). For ICP, CM was cannulated with saline-containing line, which was connected to an in-line pressure sensors (BP-102). BP and ICP were measured by using the same type of sensor. Data were collected with an iworx® IX-RA-834 (Iworx Systems Inc.) data acquisition system coupled with an iWire-BIO4 iworx® 4-Channel Biopotential Recording Module and Labscribe 4 software.

### Blood pressure lowering

Either nicardipine hydrochloride (Sigma, N7510-1G) or sodium nitroprusside (SNP) (Sigma, PHR1423-1G) was infused into the femoral vein at rates of 5 or 10 µl min^-1^, respectively. Dosages were 5.0 µg kg^-1^ min^-1^ for SNP and 0.5 µg kg^-1^ min^-1^ for nicardipine. Control groups received saline i.v. at similar rates. The i.v. infusions were 10 min for SNP and 40 min for nicardipine.

### Deep cervical lymph node ligation experiments

DcLNs were accessed through neck incision. Afferent lymphatic vessels draining into dcLNs were ligated with surgical sutures. Sham animals received otherwise similar surgery without the ligation.

### Blood collection and fluorescence analysis of serum

160 – 200 µl of blood was collected and centrifuged using a table-top centrifuge at 1100 g for 10 min and the supernatant (serum) stored at -70°C. Thawed serum samples were diluted with 1xPBS 1:4 and assayed in triplicates using a fluorometer (PerkinElmer, VICTOR3V 1420 Multilabel counter) with excitation/emission of 485 nm / 535 nm (25 nm band-pass filter). Serum without FD40 served as negative control.

### Microscopical fluorescence analysis from brain and deep cervical lymph nodes

Brain and dcLNs containing fixable 45 kDa Texas Red-conjugated ovalbumin were isolated and drop-fixed in 4% paraformaldehyde overnight at +4°C. On the following day, samples were transferred to 1xPBS with 0.02% Sodium Azide until further processing. Brains were sectioned coronally from bregma +1.8 mm to –2.4 mm into 100-µm free floating sections using a vibratome (VT1200S, Leica) and collected in 1xPBS (with 0.02% sodium azide). Every 6^th^ section was selected, and incubated in PBS containing 4′,6-diamidino-2-phenylindole (DAPI, dilution 1:1000) overnight at +4°C on a shaker. Brain slices were mounted on glass slides, covered with a cover slip, and stored at +4°C until imaging. Brain and lymph nodes containing ovalbumin-Texas Red were imaged using an AxioZoom V16 stereomicroscope (CarlZeiss) equipped with an Orca-Fusion BT sCMOS camera (Hamamatsu Systems) using Zen 3.3 pro software (CarlZeiss). The objective used was PlanNeoFluar Z 2.3x and PlanNeoFluar Z 1.0x, respectively and imaging parameters were the same for all the samples. Quantitative analysis of the acquired images was performed using the Fiji processing package of Image J2 software (Version 1.53, National Institute of Health). For perivascular influx analysis, the background was subtracted uniformly from the brain sections. The slices were manually outlined, mean pixel intensity was measured and the average value of 7 sections per animal was reported. A free hand -tool was used to make a region of interest and mean pixel intensities of deep cervical lymph nodes from each animal were measured and the average value was reported.

### Statistical analyses

Values are mean ± standard error of mean (SEM) or 95% confidence interval (CI, data shown across multiple timepoints). Unpaired t-test was used for comparison of two groups. In the case of unequal variances Welch’s correction was used. If data was not normally distributed as shown by Shapiro-Wilk test, Mann-Whitney *U* test was used. In experiments with more than two groups, ordinary two-way analysis of variance (ANOVA) followed by post hoc Šidák’s multiple comparisons testing was employed. For multiple timepoints, where each timepoint was compared between two different populations, multiple unpaired t-tests with Welch’s correction were used. Alternatively, ordinary one-way ANOVA with repeated measures and Greenhouse-Geisser correction followed by Dunnett’s multiple comparisons test was employed, where each timepoint was tested against the first timepoint after the baseline. Values for multiple timepoints are presented as fold changes instead of absolute values due to high physiological variability of baseline values in individual mice. *P* < 0.05 was considered statistically significant. All analyses were performed with GraphPad Prism (version 9.3.1, GraphPad Software, Inc; San Diego, Ca, USA).

## Declarations

### Ethics approval and consent to participate

All experimental procedures and animal care were in accordance with the Finnish and European legislation and were approved by the Finnish National Project Authorization Board (license numbers ESAVI/41363/2019 and ESAVI/2362/04.10.07/2017).

### Availability of data and materials

The datasets used and/or analyzed during the current study are available from the corresponding author on reasonable request.

## Funding

This research was funded by the Health from Science program (TERVA) of the Academy of Finland (the role of the funding; the study and collection, analysis, and interpretation of data and in writing the manuscript), the Finnish Brain Foundation (collection, analysis, and interpretation of data), Jane and Aatos Erkko foundation grant (collection, analysis, and interpretation of data).

### Authors’ contributions

LE, MK, TM, JJ and AS planned and conceived the experiments. JJ performed FD40 - injection experiments, BP measurements and data analysis. JJ and MK performed ICP measurements and data analysis. AS performed Texas Red-conjugated ovalbumin drainage and brain influx experiments, all related sample processing, imaging, and data analysis. JJ, MK, AS, LE, SM, HF and TM participated in writing the original draft and the data interpretation. LE, MK, TM and VK interpreted the results and supervised the research. All authors revised and approved the final manuscript.

## Acknowledgements

The authors acknowledge the excellent technical assistance of Riitta Jokela and Jaana Träskelin. Biocenter Oulu and Light Microscopy Core Facilities supported by Biocenter Finland and the University of Oulu, and Oulu Laboratory Animal Center, University of Oulu, are acknowledged for their specific scientific expertise and research infrastructure services.

## Abbreviations

*AD*: Alzheimer’s disease
*ANOVA*: analysis of variance
*BP*: Blood pressure
*BPL*: Blood pressure lowering
*CI*: Confidence interval
*CM*: Cisterna magna
*CNS*: Central nervous system
*CSF*: Cerebrospinal fluid
*DAPI*: 4′,6-diamidino-2-phenylindole
*dcLN*: Deep cervical lymph node
*FD40*: FITC-dextran 40 kDa
*ICP*: Intracranial pressure
*i.v.*: intravenous
*KX*: Ketamine-xylazine
*NMDA*: N-methyl-D-aspartate
*NO*: Nitric oxide
*SEM*: Standard error of mean
*SNP*: Sodium nitroprusside

## References

Ahn, J. H., Cho, H., Kim, J. H., Kim, S. H., Ham, J. S., Park, I., Suh, S. H., Hong, S. P., Song, J. H., Hong, Y. K., Jeong, Y., Park, S. H., & Koh, G. Y. (2019). Meningeal lymphatic vessels at the skull base drain cerebrospinal fluid. Nature, 572(7767), 62–66. https://doi.org/10.1038/s41586-019-1419-5

Aldea, R., Weller, R. O., Wilcock, D. M., Carare, R. O., & Richardson, G. (2019). Cerebrovascular smooth muscle cells as the drivers of intramural periarterial drainage of the brain. Frontiers in Aging Neuroscience, 11(JAN). https://doi.org/10.3389/fnagi.2019.00001

Anile, C., Zanghi, F., Bracali, A., Maira, G., & Rossi, G. F. (1981). Sodium nitroprusside and intracranial pressure. Acta Neurochirurgica, 58(3–4), 203–211. https://doi.org/10.1007/BF01407126

Antila, S., Karaman, S., Nurmi, H., Airavaara, M., Voutilainen, M. H., Mathivet, T., Chilov, D., Li, Z., Koppinen, T., Park, J. H., Fang, S., Aspelund, A., Saarma, M., Eichmann, A., Thomas, J. L., & Alitalo, K. (2017a). Development and plasticity of meningeal lymphatic vessels. Journal of Experimental Medicine, 214(12), 3645–3667. https://doi.org/10.1084/jem.20170391

Antila, S., Karaman, S., Nurmi, H., Airavaara, M., Voutilainen, M. H., Mathivet, T., Chilov, D., Li, Z., Koppinen, T., Park, J. H., Fang, S., Aspelund, A., Saarma, M., Eichmann, A., Thomas, J. L., & Alitalo, K. (2017b). Development and plasticity of meningeal lymphatic vessels. The Journal of Experimental Medicine, 214(12), 3645. https://doi.org/10.1084/JEM.20170391

Asano, Y., Koehler, R. C., Kawaguchi, T., & McPherson, R. W. (1997). Pial arteriolar constriction to α2-adrenergic agonist dexmedetomidine in the rat. American Journal of Physiology - Heart and Circulatory Physiology, *272*(6 41-6). https://doi.org/10.1152/ajpheart.1997.272.6.h2547

Aspelund, A., Antila, S., Proulx, S. T., Karlsen, T. V., Karaman, S., Detmar, M., Wiig, H., & Alitalo, K. (2015). A dural lymphatic vascular system that drains brain interstitial fluid and macromolecules. Journal of Experimental Medicine, 212(7), 991–999. https://doi.org/10.1084/jem.20142290

Attier-Zmudka, J., Sérot, J.-M., Valluy, J., Saffarini, M., Macaret, A.-S., Diouf, M., Dao, S., Douadi, Y., Malinowski, K. P., & Balédent, O. (2019). Decreased Cerebrospinal Fluid Flow Is Associated With Cognitive Deficit in Elderly Patients. Frontiers in Aging Neuroscience, 11(APR), 87. https://doi.org/10.3389/fnagi.2019.00087

Balestreri, M., Czosnyka, M., Hutchinson, P., Steiner, L. A., Hiler, M., Smielewski, P., & Pickard, J. D. (2006). Impact of Intracranial Pressure and Cerebral Perfusion Pressure on Severe Disability and Mortality After Head Injury. Neurocritical Care, 4(1), 008–013. https://doi.org/10.1385/ncc:4:1:008

Bolte, A. C., Dutta, A. B., Hurt, M. E., Smirnov, I., Kovacs, M. A., McKee, C. A., Ennerfelt, H. E., Shapiro, D., Nguyen, B. H., Frost, E. L., Lammert, C. R., Kipnis, J., & Lukens, J. R. (2020). Meningeal lymphatic dysfunction exacerbates traumatic brain injury pathogenesis. Nature Communications, 11(1), 4524. https://doi.org/10.1038/s41467-020-18113-4

Bradbury, M. W. B., Cserr, H. F., & Westrop, R. J. (1981). Drainage of cerebral interstitial fluid into deep cervical lymph of the rabbit. The American Journal of Physiology, 240(4), 329–336. https://doi.org/10.1152/AJPRENAL.1981.240.4.F329

Brady, M., Rahman, A., Combs, A., Venkatraman, C., Kasper, R. T., McQuaid, C., Kwok, W. C. E., Wood, R. W., & Deane, R. (2020). Cerebrospinal fluid drainage kinetics across the cribriform plate are reduced with aging. Fluids and Barriers of the CNS, 17(1). https://doi.org/10.1186/s12987-020-00233-0

Bukke, V. N., Archana, M., Villani, R., Romano, A. D., Wawrzyniak, A., Balawender, K., Orkisz, S., Beggiato, S., Serviddio, G., & Cassano, T. (2020). The Dual Role of Glutamatergic Neurotransmission in Alzheimer’s Disease: From Pathophysiology to Pharmacotherapy. International Journal of Molecular Sciences, 21(20), 7452. https://doi.org/10.3390/ijms21207452

Choi, Y. B., Tenneti, L., Le, D. A., Ortiz, J., Bai, G., Chen, H. S. V., & Lipton, S. A. (2000). Molecular basis of NMDA receptor-coupled ion channel modulation by S-nitrosylation. Nature Neuroscience, 3(1), 15–21. https://doi.org/10.1038/71090

Clark, W. E. le G. (1920). On the Pacchionian Bodies. Journal of Anatomy, 55(Pt 1), 40. https://www.ncbi.nlm.nih.gov/pmc/articles/PMC1262928/

Cottrell, J. E., Patel, K., Turndorf, H., & Ransohoff, J. (1978). Intracranial pressure changes induced by sodium nitroprusside in patients with intracranial mass lesions. Journal of Neurosurgery, 48(3), 329–331. https://doi.org/10.3171/jns.1978.48.3.0329

da Mesquita, S., Louveau, A., Vaccari, A., Smirnov, I., Cornelison, R. C., Kingsmore, K. M., Contarino, C., Onengut-Gumuscu, S., Farber, E., Raper, D., Viar, K. E., Powell, R. D., Baker, W., Dabhi, N., Bai, R., Cao, R., Hu, S., Rich, S. S., Munson, J. M., … Kipnis, J. (2018). Functional aspects of meningeal lymphatics in ageing and Alzheimer’s disease. Nature, 560(7717), 185–191. https://doi.org/10.1038/s41586-018-0368-8

Da Mesquita, S., Papadopoulos, Z., Dykstra, T., Brase, L., Farias, F. G., Wall, M., Jiang, H., Kodira, C. D., de Lima, K. A., Herz, J., Louveau, A., Goldman, D. H., Salvador, A. F., Onengut-Gumuscu, S., Farber, E., Dabhi, N., Kennedy, T., Milam, M. G., Baker, W., … Kipnis, J. (2021). Meningeal lymphatics affect microglia responses and anti-Aβ immunotherapy. Nature, 593(7858), 255–260. https://doi.org/10.1038/s41586-021-03489-0

das Neves, S. P., Delivanoglou, N., & da Mesquita, S. (2021). CNS-Draining Meningeal Lymphatic Vasculature: Roles, Conundrums and Future Challenges. In Frontiers in Pharmacology (Vol. 12, p. 964). Frontiers Media S.A. https://doi.org/10.3389/fphar.2021.655052

Drummond, J. C., Dao, A. v., Roth, D. M., Cheng, C. R., Atwater, B. I., Minokadeh, A., Pasco, L. C., & Patel, P. M. (2008). Effect of dexmedetomidine on cerebral blood flow velocity, cerebral metabolic rate, and carbon dioxide response in normal humans. Anesthesiology, 108(2), 225–232. https://doi.org/10.1097/01.anes.0000299576.00302.4c

Elamaa, H., Kihlström, M., Kapiainen, E., Kaakinen, M., Miinalainen, I., Ragauskas, S., Cerrada-Gimenez, M., Mering, S., Nätynki, M., & Eklund, L. (2018). Angiopoietin-4-dependent venous maturation and fluid drainage in the peripheral retina. ELife, 7. https://doi.org/10.7554/eLife.37776

Fleischman, D., Berdahl, J. P., Zaydlarova, J., Stinnett, S., Fautsch, M. P., & Allingham, R. R. (2012). Cerebrospinal Fluid Pressure Decreases with Older Age. PLoS ONE, 7(12), e52664. https://doi.org/10.1371/journal.pone.0052664

Flexner, L. B. (1933). Some Problems of the Origin, Circulation and Absorption of the Cerebrospinal Fluid. The Quarterly Review of Biology, 8(4). https://doi.org/10.1086/394447

Gaab, M., Czech, T., & Korn, A. (1985). Intracranial effects of nicardipine. British Journal of Clinical Pharmacology, 20(1 S), 67S–74S. https://doi.org/10.1111/j.1365-2125.1985.tb05145.x

Greenamyre, J. T., Maragos, W. F., Albin, R. L., Penney, J. B., & Young, A. B. (1988). Glutamate transmission and toxicity in alzheimer’s disease. Progress in Neuropsychopharmacology and Biological Psychiatry, 12(4). https://doi.org/10.1016/0278-5846(88)90102-9

Harrison, I. F., Ismail, O., Machhada, A., Colgan, N., Ohene, Y., Nahavandi, P., Ahmed, Z., Fisher, A., Meftah, S., Murray, T. K., Ottersen, O. P., Nagelhus, E. A., O’Neill, M. J., Wells, J. A., & Lythgoe, M. F. (2020). Impaired glymphatic function and clearance of tau in an Alzheimer’s disease model. Brain, 143(8), 2576–2593. https://doi.org/10.1093/brain/awaa179

Hoffmann, J., Kreutz, K. M., Csapó-Schmidt, C., Becker, N., Kunte, H., Fekonja, L. S., Jadan, A., & Wiener, E. (2019). The effect of CSF drain on the optic nerve in idiopathic intracranial hypertension. Journal of Headache and Pain, 20(1), 1–10. https://doi.org/10.1186/S10194-019-1004-1/TABLES/2

Holstein-Rønsbo, S., Gan, Y., Giannetto, M. J., Rasmussen, M. K., Sigurdsson, B., Beinlich, F. R. M., Rose, L., Untiet, V., Hablitz, L. M., Kelley, D. H., & Nedergaard, M. (2023). Glymphatic influx and clearance are accelerated by neurovascular coupling. Nature Neuroscience 2023 26:6, 26(6), 1042–1053. https://doi.org/10.1038/s41593-023-01327-2

Hsu, S. J., Zhang, C., Jeong, J., Lee, S. il, McConnell, M., Utsumi, T., & Iwakiri, Y. (2021). Enhanced Meningeal Lymphatic Drainage Ameliorates Neuroinflammation and Hepatic Encephalopathy in Cirrhotic Rats. Gastroenterology, 160(4), 1315–1329.e13. https://doi.org/10.1053/j.gastro.2020.11.036

Huotari, N., Tuunanen, J., Raitamaa, L., Raatikainen, V., Kananen, J., Helakari, H., Tuovinen, T., Järvelä, M., Kiviniemi, V., & Korhonen, V. (2022). Cardiovascular Pulsatility Increases in Visual Cortex Before Blood Oxygen Level Dependent Response During Stimulus. Frontiers in Neuroscience, 16, 836378. https://doi.org/10.3389/FNINS.2022.836378/BIBTEX

Ikeda, U., Ikeda, M., Kano, S., Kanbe, T., & Shimada, K. (1996). Effect of cilostazol, a cAMP phosphodiesterase inhibitor, on nitric oxide production by vascular smooth muscle cells. European Journal of Pharmacology, 314(1–2), 197–202. https://doi.org/10.1016/S0014-2999(96)00551-1

Iliff, J. J., Wang, M., Liao, Y., Plogg, B. A., Peng, W., Gundersen, G. A., Benveniste, H., Vates, G. E., Deane, R., Goldman, S. A., Nagelhus, E. A., & Nedergaard, M. (2012). A Paravascular Pathway Facilitates CSF Flow Through the Brain Parenchyma and the Clearance of Interstitial Solutes, Including Amyloid β. Science Translational Medicine, 4(147), 147ra111. https://doi.org/10.1126/SCITRANSLMED.3003748

Iliff, J. J., Wang, M., Zeppenfeld, D. M., Venkataraman, A., Plog, B. A., Liao, Y., Deane, R., & Nedergaard, M. (2013a). Cerebral Arterial Pulsation Drives Paravascular CSF-Interstitial Fluid Exchange in the Murine Brain. Journal of Neuroscience, 33(46), 18190–18199. https://doi.org/10.1523/JNEUROSCI.1592-13.2013

Iliff, J. J., Wang, M., Zeppenfeld, D. M., Venkataraman, A., Plog, B. A., Liao, Y., Deane, R., & Nedergaard, M. (2013b). Cerebral arterial pulsation drives paravascular CSF-interstitial fluid exchange in the murine brain. The Journal of Neuroscience : The Official Journal of the Society for Neuroscience, 33(46), 18190–18199. https://doi.org/10.1523/JNEUROSCI.1592-13.2013

Jiang, Y., Meng, L., Yan, J., Yue, H., Zhu, J., & Liu, Y. (2021). Changes of arachnoid granulations after subarachnoid hemorrhage in cynomolgus monkeys. Journal of Integrative Neuroscience, 20(2), 419–424. https://doi.org/10.31083/j.jin2002043

Johnston, M., Zakharov, A., Papaiconomou, C., Salmasi, G., & Armstrong, D. (2004). Evidence of connections between cerebrospinal fluid and nasal lymphatic vessels in humans, non-human primates and other mammalian species. Cerebrospinal Fluid Research, 1(1), 1–13. https://doi.org/10.1186/1743-8454-1-2/COMMENTS

Keil, S. A., Braun, M., O’Boyle, R., Sevao, M., Pedersen, T., Agarwal, S., Jansson, D., & Iliff, J. J. (2022). Dynamic infrared imaging of cerebrospinal fluid tracer influx into the brain. Neurophotonics, 9(03). https://doi.org/10.1117/1.nph.9.3.031915

Kellie, G. (1824). An Account of the Appearances Observed in the Dissection of Two of Three Individuals Presumed to Have Perished in the Storm of the 3d, and Whose Bodies Were Discovered in the Vicinity of Leith on the Morning of the 4th, November 1821; with Some Reflections on the Pathology of the Brain: Part I. Transactions. Medico-Chirurgical Society of Edinburgh, 1, 84. https://www-ncbi-nlm-nih-gov.pc124152.oulu.fi:9443/pmc/articles/PMC5405298/

Kim, D. J., Kasprowicz, M., Carrera, E., Castellani, G., Zweifel, C., Lavinio, A., Smielewski, P., Sutcliffe, M. P. F., Pickard, J. D., & Czosnyka, M. (2009). The monitoring of relative changes in compartmental compliances of brain. Physiological Measurement, 30(7), 647. https://doi.org/10.1088/0967-3334/30/7/009

Kitade, H., Sakitani, K., Indue, K., Masu, Y., Kawada, N., Hiramatsu, Y., Kamiyama, Y., Okumura, T., & Ito, S. (1996). Interleukin 1β markedly stimulates nitric oxide formation in the absence of other cytokines or lipopolysaccharide in primary cultured rat hepatocytes but not in Kupffer cells. Hepatology, 23(4), 797–802. https://doi.org/10.1053/jhep.1996.v23.pm0008666334

Kress, B. T., Iliff, J. J., Xia, M., Wang, M., Wei Bs, H. S., Zeppenfeld, D., Xie, L., Hongyi Kang, B. S., Xu, Q., Liew, J. A., Plog, B. A., Ding, F., PhD, R. D., & Nedergaard, M. (2014). Impairment of paravascular clearance pathways in the aging brain. Annals of Neurology, 76(6), 845–861. https://doi.org/10.1002/ANA.24271

Lahiri, S., Nezhad, M., Schlick, K. H., Rinsky, B., Rosengart, A., Mayer, S. A., & Lyden, P. D. (2018). Paradoxical cerebrovascular hemodynamic changes with nicardipine. Journal of Neurosurgery, 128(4), 1015–1019. https://doi.org/10.3171/2016.11.JNS161992

Lee, J., Jang, H., Kang, S. H., Kim, J., Kim, J. S., Kim, J. P., Kim, H. J., Seo, S. W., & Na, D. L. (2020). Cerebrospinal Fluid Biomarkers for the Diagnosis and Classification of Alzheimer’s Disease Spectrum. Journal of Korean Medical Science, 35(44). https://doi.org/10.3346/jkms.2020.35.e361

Levy, J. H. (2005). Management of systemic and pulmonary hypertension. Texas Heart Institute Journal, 32(4). https://doi.org/10.1007/978-88-470-0558-7_15

Li, Q. Q., Chen, J., Hu, P., Jia, M., Sun, J. H., Feng, H. Y., Qiao, F. C., Zang, Y. Y., Shi, Y. Y., Chen, G., Sheng, N., Xu, Y., Yang, J. J., Xu, Z., & Shi, Y. S. (2022). Enhancing GluN2A-type NMDA receptors impairs long-term synaptic plasticity and learning and memory. Molecular Psychiatry, 1–11. https://doi.org/10.1038/s41380-022-01579-7

Lilius, T. O., Blomqvist, K., Hauglund, N. L., Liu, G., Stæger, F. F., Bærentzen, S., Du, T., Ahlström, F., Backman, J. T., Kalso, E. A., Rauhala, P. v., & Nedergaard, M. (2019). Dexmedetomidine enhances glymphatic brain delivery of intrathecally administered drugs. Journal of Controlled Release, 304, 29–38. https://doi.org/10.1016/j.jconrel.2019.05.005

Liu, Z., Yang, S., Jin, X., Zhang, G., Guo, B., Chen, H., Yu, P., Sun, Y., Zhang, Z., & Wang, Y. (2017). Synthesis and biological evaluation of memantine nitrates as a potential treatment for neurodegenerative diseases. MedChemComm, 8(1), 135–147. https://doi.org/10.1039/c6md00509h

Louveau, A., Plog, B. A., Antila, S., Alitalo, K., Nedergaard, M., & Kipnis, J. (2017). Understanding the functions and relationships of the glymphatic system and meningeal lymphatics. The Journal of Clinical Investigation, 127(9), 3210–3219. https://doi.org/10.1172/JCI90603

Louveau, A., Smirnov, I., Keyes, T. J., Eccles, J. D., Rouhani, S. J., Peske, J. D., Derecki, N. C., Castle, D., Mandell, J. W., Lee, K. S., Harris, T. H., & Kipnis, J. (2015). Structural and functional features of central nervous system lymphatic vessels. Nature, 523(7560), 337–341. https://doi.org/10.1038/nature14432

Luo, F., Wu, L., Zhang, Z., Zhu, Z., Liu, Z., Guo, B., Li, N., Ju, J., Zhou, Q., Li, S., Yang, X., Mak, S., Han, Y., Sun, Y., Wang, Y., Zhang, G., & Zhang, Z. (2019). The dual-functional memantine nitrate MN-08 alleviates cerebral vasospasm and brain injury in experimental subarachnoid haemorrhage models. British Journal of Pharmacology, 176(17), bph.14763. https://doi.org/10.1111/bph.14763

Ma, Q., Ineichen, B. v., Detmar, M., & Proulx, S. T. (2017). Outflow of cerebrospinal fluid is predominantly through lymphatic vessels and is reduced in aged mice. Nature Communications, 8(1), 1434. https://doi.org/10.1038/s41467-017-01484-6

Ma, Q., Ries, M., Decker, Y., Müller, A., Riner, C., Bücker, A., Fassbender, K., Detmar, M., & Proulx, S. T. (2019). Rapid lymphatic efflux limits cerebrospinal fluid flow to the brain. Acta Neuropathologica, 137(1), 151–165. https://doi.org/10.1007/s00401-018-1916-x

Maki, T., Okamoto, Y., Carare, R. O., Hase, Y., Hattori, Y., Hawkes, C. A., Saito, S., Yamamoto, Y., Terasaki, Y., Ishibashi-Ueda, H., Taguchi, A., Takahashi, R., Miyakawa, T., Kalaria, R. N., Lo, E. H., Arai, K., & Ihara, M. (2014). Phosphodiesterase III inhibitor promotes drainage of cerebrovascular β-amyloid. Annals of Clinical and Translational Neurology, 1(8), 519–533. https://doi.org/10.1002/acn3.79

Maloveska, M., Danko, J., Petrovova, E., Kresakova, L., Vdoviakova, K., Michalicova, A., Kovac, A., Cubinkova, V., & Cizkova, D. (2018). Dynamics of Evans blue clearance from cerebrospinal fluid into meningeal lymphatic vessels and deep cervical lymph nodes. Neurological Research, 40(5), 372–380. https://doi.org/10.1080/01616412.2018.1446282

May, C., Kaye, J. A., Atack, J. R., Schapiro, M. B., Friedland, R. P., & Rapoport, S. I. (1990). Cerebrospinal fluid production is reduced in healthy aging. Neurology, 40(3 Pt 1), 500–503. https://doi.org/10.1212/WNL.40.3_PART_1.500

Mestre, H., Du, T., Sweeney, A. M., Liu, G., Samson, A. J., Peng, W., Mortensen, K. N., Stæger, F. F., Bork, P. A. R., Bashford, L., Toro, E. R., Tithof, J., Kelley, D. H., Thomas, J. H., Hjorth, P. G., Martens, E. A., Mehta, R. I., Solis, O., Blinder, P., … Nedergaard, M. (2020). Cerebrospinal fluid influx drives acute ischemic tissue swelling. Science, 367(6483). https://doi.org/10.1126/science.aaw7462

Mestre, H., Tithof, J., Du, T., Song, W., Peng, W., Sweeney, A. M., Olveda, G., Thomas, J. H., Nedergaard, M., & Kelley, D. H. (2018). Flow of cerebrospinal fluid is driven by arterial pulsations and is reduced in hypertension. Nature Communications, 9(1), 1–9. https://doi.org/10.1038/s41467-018-07318-3

Mollanji, R., Bozanovic-Sosic, R., Zakharov, A., Makarian, L., & Johnston, M. G. (2002). Blocking cerebrospinal fluid absorption through the cribriform plate increases resting intracranial pressure. American Journal of Physiology - Regulatory Integrative and Comparative Physiology, 282(6 51–6). https://doi.org/10.1152/ajpregu.00695.2001

Nagahama, Y., Fukuyama, H., Yamauchi, H., Katsumi, Y., Dong, Y., Konishi, J., & Kimura, J. (1997). Effect of nicardipine on cerebral blood flow in hypertensive patients with internal carotid artery occlusion: A PET study. Journal of Stroke and Cerebrovascular Diseases, 6(5), 325–331. https://doi.org/10.1016/S1052-3057(97)80214-5

Nishikawa, T., Omote, K., Namiki, A., & Takahashi, T. (1986). The effects of nicardipine on cerebrospinal fluid pressure in humans. Anesthesia and Analgesia, 65(5). https://doi.org/10.1213/00000539-198605000-00015

Paoletti, P., Bellone, C., & Zhou, Q. (2013). NMDA receptor subunit diversity: Impact on receptor properties, synaptic plasticity and disease. In Nature Reviews Neuroscience (Vol. 14, Issue 6, pp. 383–400). Nature Publishing Group. https://doi.org/10.1038/nrn3504

Patel, T. K., Habimana-Griffin, L., Gao, X., Xu, B., Achilefu, S., Alitalo, K., McKee, C. A., Sheehan, P. W., Musiek, E. S., Xiong, C., Coble, D., & Holtzman, D. M. (2019). Dural lymphatics regulate clearance of extracellular tau from the CNS. Molecular Neurodegeneration, 14(1), 11. https://doi.org/10.1186/s13024-019-0312-x

Pedler, M. G., Petrash, J. M., & Subramanian, P. S. (2021). Prostaglandin analog effects on cerebrospinal fluid reabsorption via nasal mucosa. PLOS ONE, 16(12), e0248545. https://doi.org/10.1371/journal.pone.0248545

Plog, B. A., Mestre, H., Olveda, G. E., Sweeney, A. M., Kenney, H. M., Cove, A., Dholakia, K. Y., Tithof, J., Nevins, T. D., Lundgaard, I., Du, T., Kelley, D. H., & Nedergaard, M. (2018). Transcranial optical imaging reveals a pathway for optimizing the delivery of immunotherapeutics to the brain. JCI Insight, 3(20). https://doi.org/10.1172/jci.insight.120922

Potts, D. G., & Deonarine, V. (1973). Effect of positional changes and jugular vein compression on the pressure gradient across the arachnoid villi and granulations of the dog. Journal of Neurosurgery, 38(6), 722–728. https://doi.org/10.3171/jns.1973.38.6.0722

Proulx, S. T. (2021). Cerebrospinal fluid outflow: a review of the historical and contemporary evidence for arachnoid villi, perineural routes, and dural lymphatics. In Cellular and Molecular Life Sciences (pp. 1–29). Springer Science and Business Media Deutschland GmbH. https://doi.org/10.1007/s00018-020-03706-5

Rubenstein, E. (1998). Relationship of senescence of cerebrospinal fluid circulatory system to dementias of the aged. In Lancet (Vol. 351, Issue 9098, pp. 283–285). Lancet Publishing Group. https://doi.org/10.1016/S0140-6736(97)09234-9

Sahay, N., Bhadani, U., Guha, S., Himanshu, A., Sinha, C., Bara, M., Sahay, A., Ranjan, A., & Singh, P. (2018). Effect of dexmedetomidine on intracranial pressures during laparoscopic surgery: A randomized, placebo-controlled trial. Journal of Anaesthesiology Clinical Pharmacology, 34(3), 341–346. https://doi.org/10.4103/joacp.JOACP_171_17

Silver, I., Kim, C., Mollanji, R., & Johnston, M. (2002). Cerebrospinal fluid outflow resistance in sheep: impact of blocking cerebrospinal fluid transport through the cribriform plate. Neuropathology and Applied Neurobiology, 28(1), 67–74. https://doi.org/10.1046/j.1365-2990.2002.00373.x

Simon, M. J., & Iliff, J. J. (2016). Regulation of cerebrospinal fluid (CSF) flow in neurodegenerative, neurovascular and neuroinflammatory disease. Biochimica et Biophysica Acta, 1862(3), 442–451. https://doi.org/10.1016/j.bbadis.2015.10.014

Song, E., Mao, T., Dong, H., Boisserand, L. S. B., Antila, S., Bosenberg, M., Alitalo, K., Thomas, J. L., & Iwasaki, A. (2020). VEGF-C-driven lymphatic drainage enables immunosurveillance of brain tumours. Nature, 577(7792), 689–694. https://doi.org/10.1038/s41586-019-1912-x

Stoquart-ElSankari, S., Balédent, O., Gondry-Jouet, C., Makki, M., Godefroy, O., & Meyer, M. E. (2007). Aging effects on cerebral blood and cerebrospinal fluid flows. Journal of Cerebral Blood Flow and Metabolism, 27(9), 1563–1572. https://doi.org/10.1038/sj.jcbfm.9600462

Takenaka, T., & Handa, J. (1979). Cerebrovascular effects of YC-93, a new vasodilator, in dogs, monkeys and human patients. International Journal of Clinical Pharmacology Therapy and Toxicology, 17(1), 1–11.

Tryambake, D., He, J., Firbank, M. J., O’Brien, J. T., Blamire, A. M., & Ford, G. A. (2013). Intensive blood pressure lowering increases cerebral blood flow in older subjects with hypertension. Hypertension, 61(6),1309–1315. https://doi.org/10.1161/HYPERTENSIONAHA.112.200972

Turliuc, D. M., Sava, A., Cucu, A. I., Turliuc, Ş., Dumitrescu Costea, A.-M., & Florida, C. (2016). Cribriform Plate and Galen’ s Cribrum romanum. Rom J Anat, XV(1), 123–126.

van Veluw, S. J., Hou, S. S., Calvo-Rodriguez, M., Arbel-Ornath, M., Snyder, A. C., Frosch, M. P., Greenberg, S. M., & Bacskai, B. J. (2020). Vasomotion as a Driving Force for Paravascular Clearance in the Awake Mouse Brain. Neuron, 105(3), 549–561.e5. https://doi.org/10.1016/j.neuron.2019.10.033

Williamson, J. D., Pajewski, N. M., Auchus, A. P., Bryan, R. N., Chelune, G., Cheung, A. K., Cleveland, M. L., Coker, L. H., Crowe, M. G., Cushman, W. C., Cutler, J. A., Davatzikos, C., Desiderio, L., Erus, G., Fine, L. J., Gaussoin, S. A., Harris, D., Hsieh, M. K., Johnson, K. C., … Wright, C. B. (2019). Effect of Intensive vs Standard Blood Pressure Control on Probable Dementia: A Randomized Clinical Trial. JAMA, 321(6), 553–561. https://doi.org/10.1001/JAMA.2018.21442

Wilson, M. H. (2016). Monro-Kellie 2.0: The dynamic vascular and venous pathophysiological components of intracranial pressure. Journal of Cerebral Blood Flow and Metabolism, 36(8), 1338–1350. https://doi.org/10.1177/0271678X16648711

Wolfsgruber, S., Polcher, A., Koppara, A., Kleineidam, L., Frölich, L., Peters, O., Hüll, M., Rüther, E., Wiltfang, J., Maier, W., Kornhuber, J., Lewczuk, P., Jessen, F., & Wagner, M. (2017). Cerebrospinal Fluid Biomarkers and Clinical Progression in Patients with Subjective Cognitive Decline and Mild Cognitive Impairment. Journal of Alzheimer’s Disease, 58(3), 939–950. https://doi.org/10.3233/JAD-161252

Wood, A. M., Lequin, M. H., Philippens, M. M., Seravalli, E., Plasschaert, S. L., van den Heuvel-Eibrink, M. M., & Janssens, G. O. (2019). MRI-guided definition of cerebrospinal fluid distribution around cranial and sacral nerves: implications for brain tumors and craniospinal irradiation. Acta Oncologica, 58(12), 1740–1744. https://doi.org/10.1080/0284186X.2019.1667023

Xavier, A. L. R., Hauglund, N. L., von Holstein-Rathlou, S., Li, Q., Sanggaard, S., Lou, N., Lundgaard, I., & Nedergaard, M. (2018). Cannula implantation into the cisterna magna of rodents. Journal of Visualized Experiments, 2018(135). https://doi.org/10.3791/57378

Xiang, T., Feng, D., Zhang, X., Chen, Y., Wang, H., Liu, X., Gong, Z., Yuan, J., Liu, M., Sha, Z., Lv, C., Jiang, W., Nie, M., Fan, Y., Wu, D., Dong, S., Feng, J., Ponomarev, E. D., Zhang, J., & Jiang, R. (2022). Effects of increased intracranial pressure on cerebrospinal fluid influx, cerebral vascular hemodynamic indexes, and cerebrospinal fluid lymphatic efflux. Journal of Cerebral Blood Flow and Metabolism. https://doi.org/10.1177/0271678X221119855

Xie, L., Kang, H., Xu, Q., Chen, M. J., Liao, Y., Thiyagarajan, M., O’Donnell, J., Christensen, D. J., Nicholson, C., Iliff, J. J., Takano, T., Deane, R., & Nedergaard, M. (2013). Sleep drives metabolite clearance from the adult brain. Science (New York, N.Y.), 342(6156), 373–377. https://doi.org/10.1126/science.1241224

Yasuda, K., Sakuma, M., & Tanabe, T. (1985). Hemodynamic effect of cilostazol on increasing peripheral blood flow in arteriosclerosis obliterans. Arzneimittel-Forschung/Drug Research, 35(7 A), 1198–1200.

Yeung, J. H. Y., Walby, J. L., Palpagama, T. H., Turner, C., Waldvogel, H. J., Faull, R. L. M., & Kwakowsky, A. (2021). Glutamatergic receptor expression changes in the Alzheimer’s disease hippocampus and entorhinal cortex. Brain Pathology, 31(6), e13005. https://doi.org/10.1111/bpa.13005

Zakharov, A., Papaiconomou, C., Koh, L., Djenic, J., Bozanovic-Sosic, R., & Johnston, M. (2004). Integrating the roles of extracranial lymphatics and intracranial veins in cerebrospinal fluid absorption in sheep. Microvascular Research, *67*(1), 96–104. https://doi.org/10.1016/j.mvr.2003.08.004

